# Multi-omics analyses reveal environmentally adaptive cell-state remodeling during human iPSC biomanufacturing

**DOI:** 10.64898/2026.05.26.727850

**Authors:** James Colter, Tiffany Dang, Emilie J. Gysel, Daniel Young, Antoine Dufour, Ian Lewis, Kartikeya Murari, Michael Scott Kallos

**Affiliations:** Pharmaceutical Production Research Facility (PPRF), University of Calgary, 2500 University Drive NW, Calgary, AB T2N 1N4, Canada; Integrated Circuits and Optical Imaging Laboratory (ICOI), University of Calgary, 2500 University Drive NW, Calgary, AB T2N 1N4, Canada; McCaig Institute for Bone and Joint Health, University of Calgary, 3280 Hospital Drive NW, Calgary, AB, T2N 4Z6, Canada; Alberta Children’s Hospital Research Institute, University of Calgary, 3330 Hospital Dr. NW, Calgary, AB, T2N 4N1, Canada; Hotchkiss Brain Institute, University of Calgary, 3330 Hospital Dr. NW, Calgary, AB, T2N 4N1, Canada; Department of Biomedical Engineering, Schulich School of Engineering, University of Calgary, 2500 University Drive NW, Calgary, AB T2N 1N4, Canada; Department of Physiology and Pharmacology, Cumming School of Medicine, University of Calgary, 3330 Hospital Drive NW, Calgary, AB T2N 1N4, Canada; Department of Biological Sciences, Faculty of Science, University of Calgary, 2500 University Drive NW, Calgary, AB T2N 1N4, Canada; Faculty of Veterinary Medicine, University of Calgary, 3330 Hospital Drive NW, Calgary, AB T2N 1N4, Canada; Snyder Institute for Chronic Diseases, University of Calgary, 3330 Hospital Dr. NW, Calgary, AB, T2N 4N1, Canada

**Author notes:** (corresponding author), 3330 Hospital Drive NW, T2N 1N4, Calgary, AB, Canada. 2500 University Drive NW, T2N 1N4, Calgary, AB, Canada. 3330 Hospital Drive NW, T2N 1N4, Calgary, AB, Canada. 2500 University Drive NW, T2N 1N4, Calgary, AB, Canada.

## Abstract

Human induced pluripotent stem cells can retain conventional markers of pluripotency while acquiring environmentally dependent molecular states that may influence their developmental competence. Yet, how biomanufacturing conditions reorganize the interconnected networks governing cell state remains poorly characterized. Here, we first examined compensation of hiPSCs to static aggregation and dynamic agitation by proteomics. We then examined hiPSCs during single-passage culture in stirred-suspension bioprocesses under varying oxygen and agitation conditions. Intracellular metabolomic and transcriptomic profiling revealed distinct responses across culture configuration, hydrodynamic exposure, and oxygenation. Transition into dynamic culture broadly remodeled mitochondrial organization, carbon allocation, mechanotransduction, proteostasis, and developmental regulation. Increasing agitation produced a persistent response involving cellular architecture, growth-factor signaling, genome maintenance, and Epiblast- and lineage-aligned programs, whereas oxygenation elicited a smaller and more transient metabolic and transcriptional response. Through comparison with a human gastrulation reference, we show that these adaptations intersect with natural developmental programs without reproducing coherent epiblast state or lineage commitment. These findings demonstrate that bioprocess conditions can preserve core pluripotent identity while remodeling broader molecular states associated with developmental responsiveness, providing perspective on evaluating and optimizing hiPSC quality in biomanufacturing beyond restricted marker panels.

## Introduction

Human induced pluripotent stem cells (hiPSCs) have emerged as a versatile cell source for regenerative medicine, owing to their capacity for clonal expansion and their ability to differentiate into primary germ layer derivatives [1–5]. Decades of scientific progress and significant economic investment have enabled early-phase clinical trials exploring hiPSC-derived therapeutics, alongside rapid developments in technologies supporting complex biomanufacturing pipelines [6, 7]. Despite this progress, the efficacy of stem-cell–derived therapeutics remains limited, with substantial challenges hindering clinical translation.

While efforts to mitigate genetic instability during hiPSC biomanufacturing have reduced safety risks, these solutions address only a fraction of biological complexity [8, 9]. Conventional induction and maintenance strategies do not fully reconstruct the natural embryonic epiblast state across epigenetic, transcriptional, and metabolic domains [10]. Residual somatic signatures from the donor cell population persist due to incomplete epigenetic remodeling [11], and the impacts of artificial niche dynamics during expansion remain poorly understood [12, 13]. Expansion protocols are designed to maintain a restricted set of markers for pluripotent phenotype, though *in vitro* hiPSC populations have been shown to reinforce cell-state features in response to the environment that are not observed *in vivo* [14, 15]. It is perhaps no surprise then that therapeutic derivatives generated at scale frequently exhibit only partial fidelity to the phenotypic and functional characteristics of their natural counterparts [16, 17].

Given that clinical applications require expansion of small hiPSC populations into millions or billions of cells, we sought to characterize how transcriptional, metabolic, and proteomic states respond to environmental perturbation within artificial culture niches. Previous work has shown that oxygen concentration and agitation rate influence pathways central to potency, epigenetic regulation, aggregate morphology, and proliferation. For instance, TGF-β, LIF, Wnt, and PI3K/AKT signaling are sensitive to shear and hydrodynamic stress, while MAPK/HIF activity modulates PI3K/AKT under hypoxic conditions [18–20]. Although individual pathways have been studied extensively, we hypothesized that examining networks of transcriptional, translational, and metabolic regulation relative to patterns observed in early developmental trajectories would better reveal how bioprocess conditions shape cell state.

Here, we examined complementary datasets addressing related levels of hiPSC bioprocess response. Proteomic profiling was used to characterize monolayer, static aggregate, and dynamically agitated culture, as well as agitation intensity within stirred suspension. In a separate longitudinal dynamic-culture experiment, transcriptomic and intracellular metabolomic profiling were done to assess time-dependent responses to oxygen and agitation. We further compared culture-responsive transcriptional patterns with external pluripotent and early embryonic reference datasets to assess whether the observed responses intersected with cell state signatures to suggest propagation along downstream lineages. Together, this work characterizes condition-dependent variation during hiPSC expansion, interrogates experimental observations against developmental and pathway-level interpretations, and explores perspective on the propagation of cell phenotype within biomanufacturing environments.

## Results

### Culture Platform and Agitation Rate Coordinate Phenotypic, Structural, and Metabolic Adaptation

We first asked whether common hiPSC expansion platforms establish unique pathway activity distinguishable by proteomics. Proteomic profiles were compared among static monolayer cultures, static suspended aggregates, and dynamically suspended aggregates. The three pairwise experiments were co-analyzed to identify unique translational activity associated with each platform (Figure 1a, Supplementary Figure 1a).

**Figure 1:**
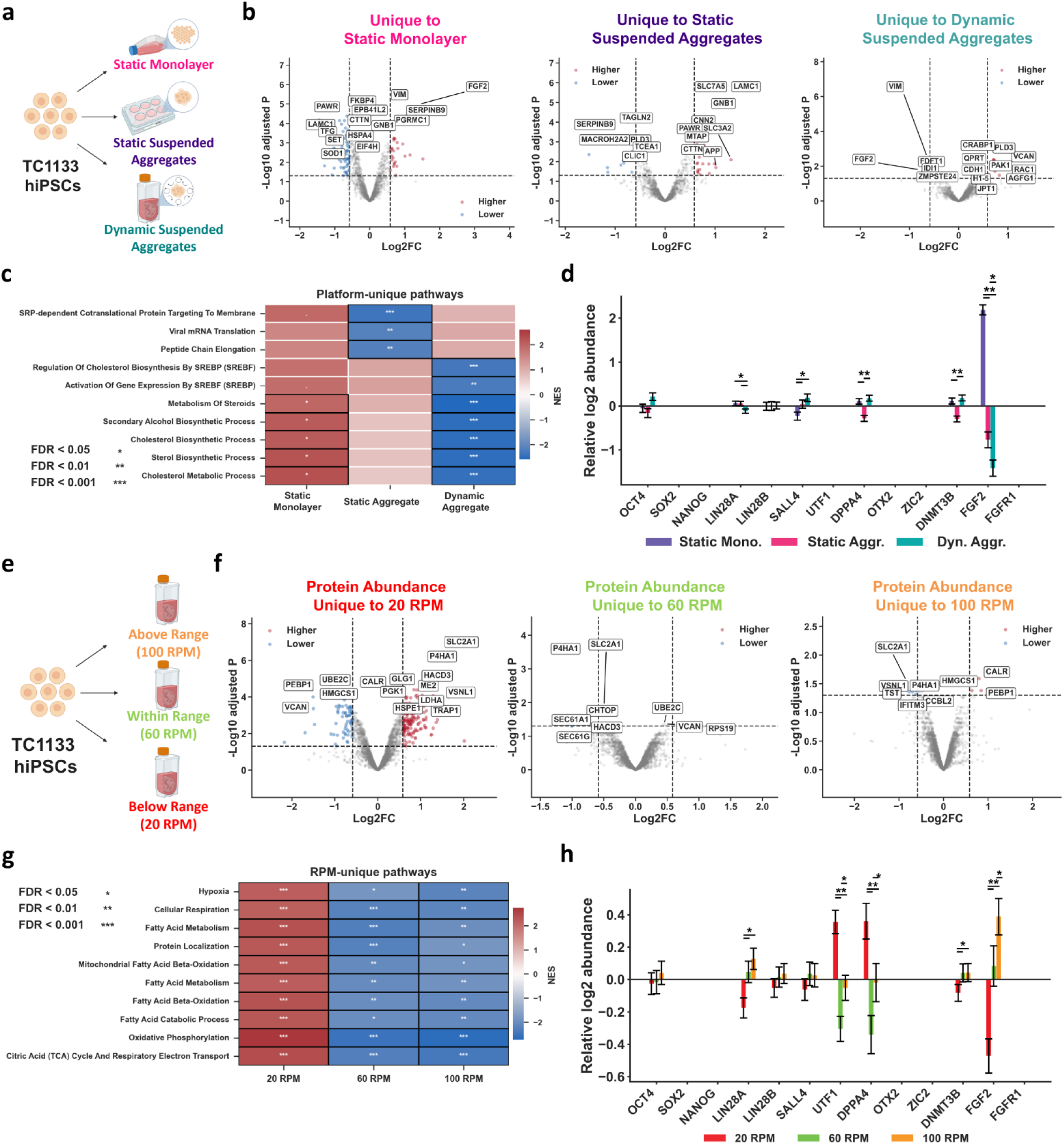
(a) Schematic representation for hiPSCs cultured on static monolayers, as static aggregates, or dynamic aggregates in stirred suspension. (b) Differential protein abundance unique within each platform. (c) Pathway enrichment for differentially abundant proteins by platform. (d) Relative abundances of selected proteins are shown by platform. (e) Schematic representation for hiPSCs cultured in stirred suspension at 20 RPM, 60 RPM, or 100 RPM. (f) Differential protein abundance unique within each dynamic agitation rate. (g) Pathway enrichment for differentially abundant proteins by dynamic agitation rate. (h) Relative abundances of selected proteins are shown by dynamic agitation rate. *Adjusted p-value or FDR < 0.05, ** adjusted p-value or FDR < 0.01, *** adjusted p-value or FDR < 0.001.

Static monolayer cultures exhibited the largest platform-specific program, whereas static and dynamic aggregates showed progressively fewer uniquely altered proteins (Supplementary Figure 1d). However, implicated phenotype genes including DPPA4, DNMT3B, and FGF2 showed significant differences in abundance, even without measurable difference in core pluripotency marker expression. Further, static suspended aggregates showed lower relative enrichment of co-translational protein targeting, peptide-chain elongation, and translation-associated programs (Figure 1b, 1c) [21]. In contrast, dynamically suspended aggregates showed lower relative enrichment of sterol and cholesterol biosynthetic programs, enriched in static monolayer culture. These findings suggest that culture configuration modulates biosynthetic and metabolic resource allocation, with subtle change in key growth-factor utilization.

We next tested whether difference in hydrodynamic exposure could further reorganize proteomic determinants of phenotypic or metabolic state (Figure 1d). Cultures maintained at 20, 60, and 100 RPM exhibited distinct protein-abundance profiles, with 20 RPM showing the largest unique response (Supplementary Figure 1e). This is expected given that such low shear stress is insufficient to support proliferation of the cell population, though viability is maintained (Supplementary Figure 1b, 1c).

Proteomic response included coordinated differences associated with cellular organization and transport, together with pronounced enrichment of hypoxia-associated, respiratory, fatty-acid metabolic, oxidative-phosphorylation, and TCA-cycle programs (Figure 1e, 1f) [22]. Agitation also modified key genes surrounding the pluripotency network (Supplementary Figure 1g). OCT4 remained comparatively stable but LIN28A, UTF1, DPPA4, and FGF2 differed among agitation rates. This supported our hypothesis that preservation of core factors alone is insufficient to conclude that dynamically expanded cells occupy equivalent phenotypic states, and that these may have broader implications.

### Environmental Perturbation Differentially Shapes Metabolic Signatures in Dynamic Culture

Following our findings of altered phenotype at the translational level, we utilized a second cell-line in our established scale-down protocol to study hiPSC maintenance at transcriptional and metabolic resolution [23]. This approach permitted us to carry out longitudinal multi omics experiments with appropriate biological and technical replicates, albeit under differing agitation rates to support near-equivalent shear stress. We were primarily interested in whether we could link transcriptional, proteomic, and metabolic signatures given our prior observations of distinct extracellular metabolic signatures [24]. We assessed cultures at day 0 following pre-aggregate formation and at days 2 and 4 of dynamic culture, pooling 2 technical replicates (t.r.) within three independent biological replicates (b.r.) (Figure 2a, 2b). Samples were taken for bulk intracellular metabolomics and RNA sequencing (see materials and methods for complete protocols). Representative morphology for cultured hiPSCs at each stage of the protocol is shown in figure 2c. Exploratory heatmaps of observed organization of metabolites across conditions and timepoints are shown in extended figure 2a-2d.

**Figure 2:**
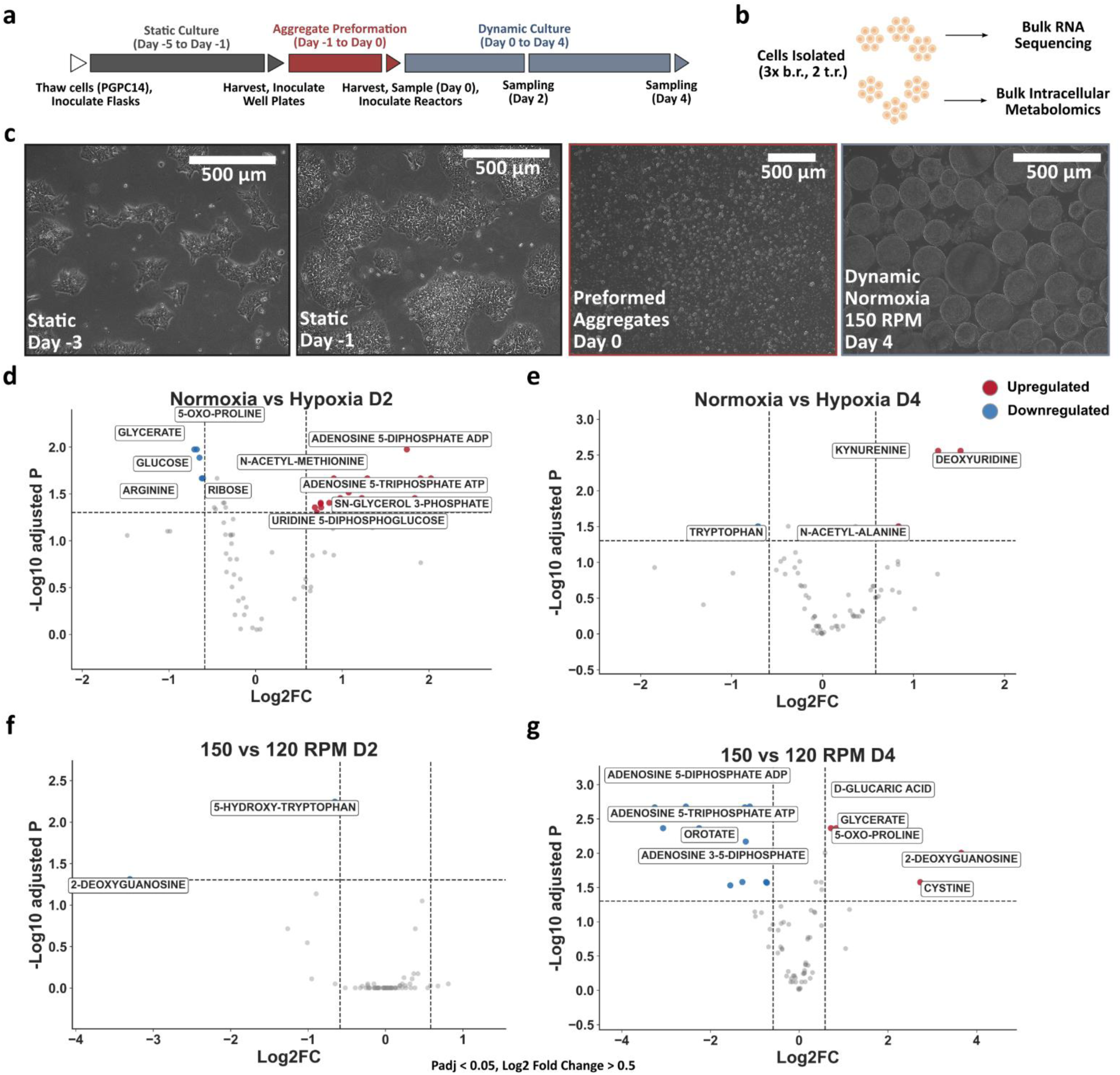
(a) Culture workflow diagram in scale-down hiPSC cultures. (b) Post-sampling replicate distributions into metabolomics and transcriptomics workflows. (c) Morphology under static, aggregate preformation, and dynamic cultures. Statistically significant differential accumulation is shown under normoxia vs. hypoxia at (d) day 2 and (e) day 4, and under 150 RPM vs. 120 RPM at (f) day 2 and (g) day 4. Dotted lines represent thresholds of Padj < 0.05, Log2FC > 0.5.

Oxygenation produced a pronounced but temporally dynamic intracellular metabolic response. At day 2, normoxia relative to hypoxia was associated with higher ADP, ATP, N-acetyl-methionine, sn-glycerol 3-phosphate, and uridine 5-diphosphoglucose, together with lower glycerate, 5-oxo-proline, glucose, arginine, and ribose (Figure 2d). These metabolites are adenylate and nucleotide-sugar pools, amino-acid-associated metabolites, glycerol-phosphate intermediates, and central carbon intermediates. By day 4, the oxygen-dependent signature had narrowed and shifted toward a more selective set of metabolites. Kynurenine, deoxyuridine, and N-acetyl-alanine were higher under normoxia, whereas tryptophan was lower (Figure 2e). The opposing behavior of tryptophan and kynurenine is consistent with altered partitioning of tryptophan-associated metabolism, while the deoxyuridine and N-acetyl-alanine responses indicate accompanying changes in pyrimidine-associated and amino-acid acetylation states [25, 26]. Divergence in these features suggests tryptophan metabolism, nucleotide homeostasis, and amino-acid acetylation as principal metabolic axes distinguishing acute normoxic and hypoxic hiPSC culture.

Agitation rate also produced time-dependent intracellular metabolic differences, although the affected metabolites differed substantially between days 2 and 4. At day 2, increasing agitation from 120 to 150 RPM was associated with lower 2-deoxyguanosine and 5-hydroxytryptophan (Figure 2f), indicating an early and relatively restricted response involving nucleoside and tryptophan-associated metabolism. By day 4, the 150 versus 120 RPM comparison showed a broader and bidirectional signature. ADP, ATP, orotate, and adenosine 3,5-diphosphate were lower at 150 RPM, whereas D-glucaric acid, glycerate, 5-oxo-proline, 2-deoxyguanosine, and cystine were higher (Figure 2g). the day 4 profile is consistent with coordinated differences in adenylate and nucleotide-associated metabolites, organic-acid and central carbon intermediates, and metabolites linked to redox and amino-acid homeostasis [27, 28].

### Environmental Perturbation Differentially Shape the hiPSC Transcriptional Landscape

To distinguish adaptation to dynamic culture from responses to oxygenation and agitation, transcriptional profiles were evaluated across three complementary contrasts: dynamic culture relative to post-aggregate initial conditions, normoxia relative to hypoxia, and 150 RPM relative to 120 RPM. Analyses were also stratified by day to determine whether transcriptional responses emerged transiently, persisted across culture, or became reinforced over time.

The three perturbations produced markedly different response magnitudes. Transition from initial aggregates into dynamic culture generated extensive transcriptional remodeling at both day 2 and day 4, indicating establishment of a temporally persistent, adapted state. Oxygen availability produced a focused response at day 2 but no genes passing statistical threshold at day 4. By contrast, the difference between 150 and 120 RPM was sparse at day 2 and remained limited at day 4, although day 4 revealed a somewhat broader collection of isolated agitation-associated genes and one significant enrichment result. Importantly, all observed differential activity occurred without significant changes in core pluripotency transcript abundance (OCT4, SOX2, NANOG).

### Dynamic Culture Establishes a Temporally Persistent Metabolically and Structurally Adapted State

Dynamic culture broadly altered the transcriptional state of hiPSCs relative to post-formed aggregates at both day 2 and day 4 (Figure 3a, 3b). The response involved coordinated changes in mitochondrial organization, lipid and sterol metabolism, ciliary trafficking, cell architecture, nutrient handling, transcriptional regulation, and cell-cycle activity.

**Figure 3:**
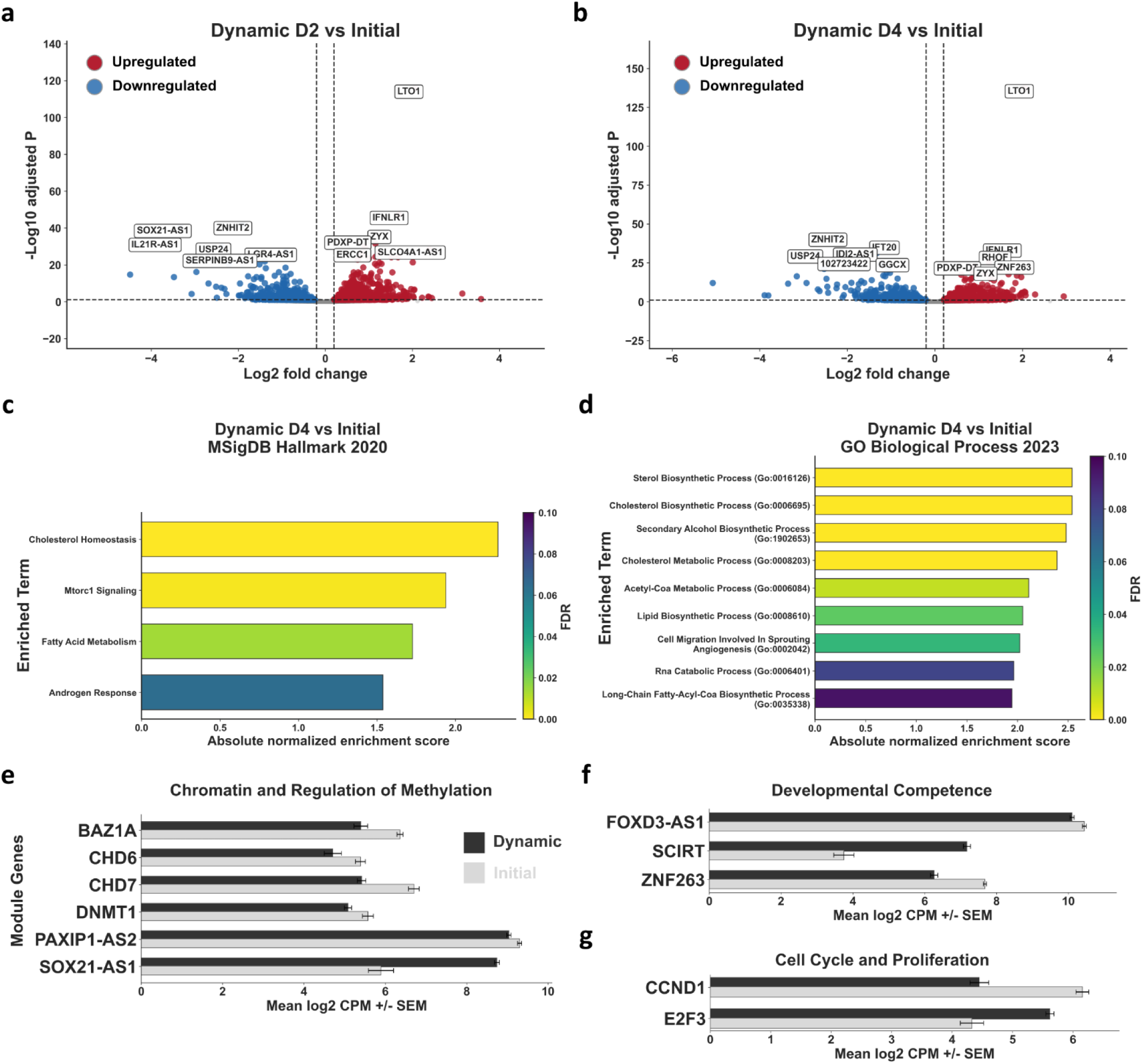
Differential gene expression at (a) day 2 and (b) day 4 following dynamic culture, pooled across oxygenation and agitation conditions. (c) Enriched MSigDB terms by day 4 under dynamic culture relative to initial, for (c) hallmark and (d) gene ontology biological processes. (e) Selected levels of expression for selected significant genes of interest across chromatin, regulation of methylation, developmental competence, cell cycle and proliferation (Padj < 0.05, Log2FC > 0.2).

Mitochondrial remodeling was not consistent with a global increase in oxidative metabolism. Dynamic culture altered genes involved in mitochondrial organization, respiratory-complex assembly, genome maintenance, and quality control [29, 30]. Enrichment analysis at day 4 independently identified cholesterol homeostasis, fatty-acid metabolism, sterol biosynthesis, cholesterol biosynthesis, cholesterol metabolism, acetyl-CoA metabolism, and long-chain fatty-acyl-CoA biosynthesis (Figure 3c, 3d).

COX11, COX15, DNM1L, HSPD1, PRIMPOL, TXNRD2, and UQCC3 changed across the dynamic transition, with the direction and magnitude varying by gene and timepoint (Supplementary Figure 3a – 3f). Dynamic culture also strongly altered carbon and lipid allocation. HK2 and other nutrient-response genes changed alongside broad repression of sterol-, cholesterol-, and fatty-acid-associated genes, including ACLY, HMGCS1, HMGCR, MVK, IDI1, SQLE, SC5D, DHCR24, FASN, SCD, and ACSL3. The dynamic transition additionally remodeled ciliary and structural programs. CFAP410, CFAP58, EVC, IFT20, IFT22, IFT74, and KIF7 identified changes in ciliary trafficking and structural organization. Changes in PTK2, PXN, RHOF, SSPN, and ZYX implicated focal adhesion, cytoskeletal organization, and force-responsive signaling. Epithelial polarity and matrix-associated genes, including ADAMTSL4, AJM1, CDHR2, CLDN22, CRB2, EDIL3, KRT8, PCDHA8, PCDHA11, PLEKHA7, PTK2, and ZYX, further demonstrated extensive reorganization of cell architecture and intercellular organization. Nutrient and energetic adaptation extended beyond lipid metabolism. Changes in ACOT2, APOE, DGAT1, HK2, IRS2, KMO, NAMPT, SLC7A11, and TFRC linked dynamic culture to altered nutrient sensing, redox support, iron handling, and metabolic resource allocation. Increased KMO provides a specific transcriptional connection to the accompanying intracellular kynurenine phenotype.

Importantly, dynamic adaptation was accompanied by changes in regulatory state. Chromatin-associated genes including BAZ1A, CHD6, CHD7, DNMT1, PAXIP1-AS2, and SOX21-AS1 remained differentially expressed (Figure 3e). FOXD3-AS1, ZNF263, and SCIRT further identified remodeling of developmental-regulatory programs (Figure 3f), while CCND1 and E2F3 supported altered cell-cycle regulation (Figure 3g). These changes indicate that dynamic culture establishes a distinct adapted hiPSC state in which metabolic and structural remodeling occurs alongside selective reconfiguration of developmental and proliferative programs [31, 32].

### Oxygenation elicits a focused and transient compensatory response

Normoxia relative to hypoxia produced a significant but comparatively restricted transcriptional response at day 2 (Figure 4a). By day 4, no genes passed the applied significance and effect-size thresholds, indicating resolution of the detectable global oxygen-associated response rather than progressive divergence between oxygen conditions (Figure 4b).

**Figure 4:**
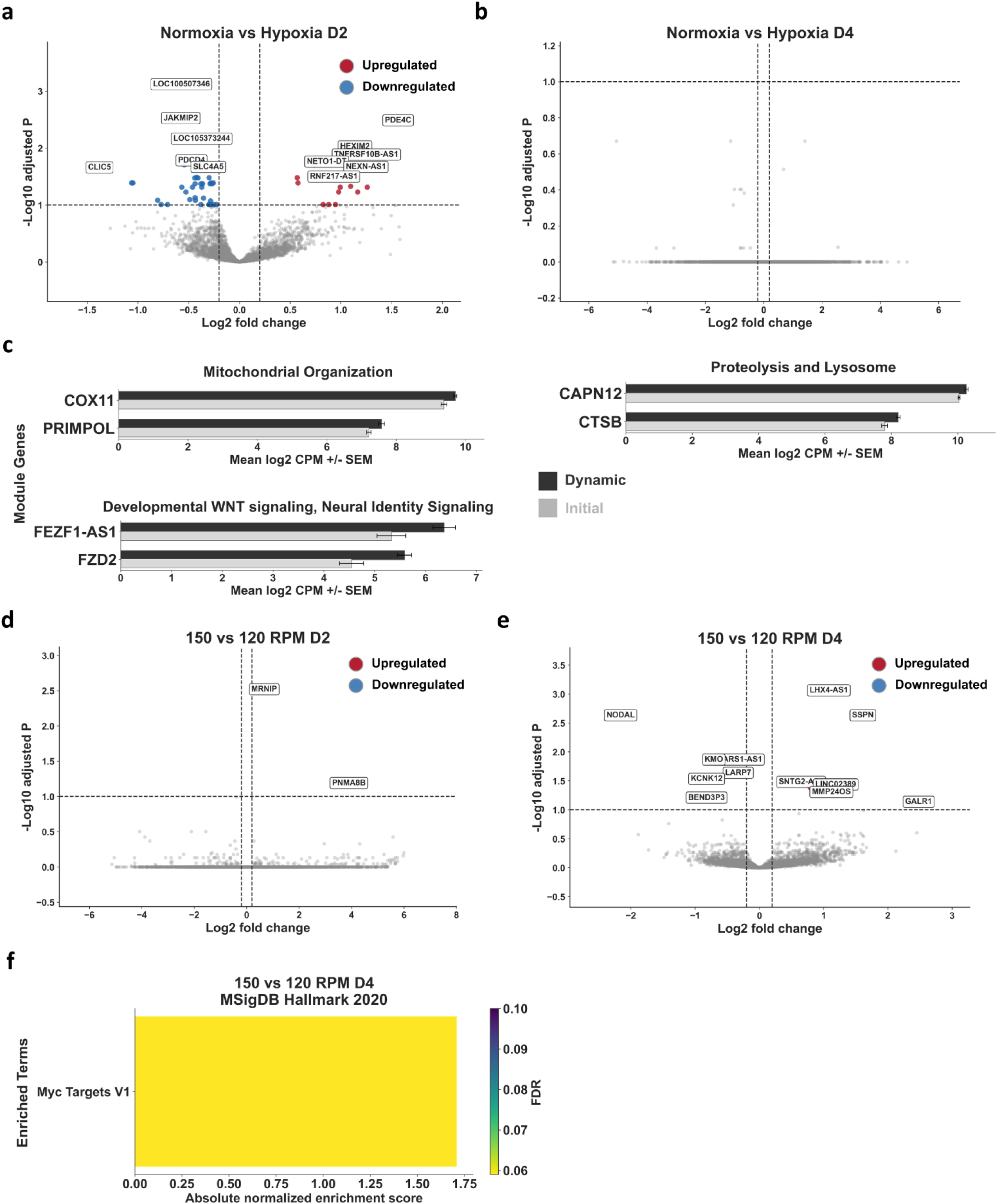
Differential gene expression at (a) day 2 and (b) day 4 under normoxia relative hypoxia, pooled across agitation conditions. (c) Selected levels of expression for selected significant genes of interest across mitochondrial organization, proteolysis, lysosome, and developmental Wnt (Padj < 0.05, Log2FC > 0.2). Differential gene expression at (d) day 2 and (e) day 4 for 150 RPM vs. 120 RPM, pooled across oxygenation conditions. (f) Sole hallmark enrichment of Myc Targets V1 at day 4 under 150 RPM vs. 120 RPM.

The day-2 oxygen response did not primarily consist of a broad canonical HIF or glycolytic program. Significant genes implicated several narrow processes. COX11 and PRIMPOL connected oxygen exposure to mitochondrial assembly and genome maintenance, whereas CAPN12 and CTSB implicated targeted proteolytic or lysosomal remodeling [33, 34]. FZD2 and FEZF1-AS1 identified a developmental WNT and neural-identity-associated component [35]. Additional significant day-2 genes included SCLT1, ADM5, and genes associated with membrane organization, trafficking, and signaling. No coherent hallmark pathways were discovered across differential genes at day 2.

### Agitation rate produces a sparse and temporally heterogeneous transcriptional response

The direct comparison between 150 and 120 RPM produced a substantially smaller transcriptional response than the transition into dynamic culture. At day 2, the response was highly restricted, with MRNIP and PNMA8B representing the clearest moderate-effect significant signals (Figure 4d). This limited gene set did not support a coherent differential gene module at day 2.

The day-4 comparison contained a broader but still limited set of agitation-associated genes (Figure 4e). Significant or prominently labeled genes included NODAL, KCNK12, LHX4-AS1, SSPN, MMP24OS, SNTG2-AS1, LINC02389, LARP7, BEND3P3, and GALR1. The mixture of coding genes, antisense transcripts, long noncoding RNAs, and isolated regulatory signals suggests that the day-4 response was heterogeneous.

Nevertheless, several genes provide biologically interpretable leads. SSPN is consistent with altered membrane-cytoskeletal organization, while KCNK12 implicates membrane excitability or ion-channel regulation [33–35]. NODAL suggests that agitation may also intersect with developmental signaling [36]. These signals remain gene-level observations, however, because the significant agitation-associated DEGs did not form a sufficiently broad shared module across days 2 and 4.

Gene-set enrichment at day 4 identified only MYC Targets V1 at the relaxed FDR threshold (Figure 4f). This isolated enrichment suggests a limited effect on a MYC-associated transcriptional program but does not support broad agitation-driven reprogramming [37]. Overall, the 150-versus-120 RPM comparison is most consistent with a sparse and time-dependent modulation layered onto a much stronger state transition induced by dynamic culture itself.

### Environmental Adaptation Reinforces Epiblast- and Lineage-Aligned Transcriptional Activity

To assess the developmental implications of adaptation to culture, we compared experimental transcriptional responses with an in vivo reference atlas of human gastrulation. Using annotations reported by Tyser et al., we calculated differential expression between gastrulating Epiblast, downstream, and extraembryonic cell types (Figure 5a) [38]. This reference was selected because the culture conditions used in the present study have been established as maintaining hiPSCs in a primed pluripotent state resembling post-implantation Epiblast [39].

**Figure 5:**
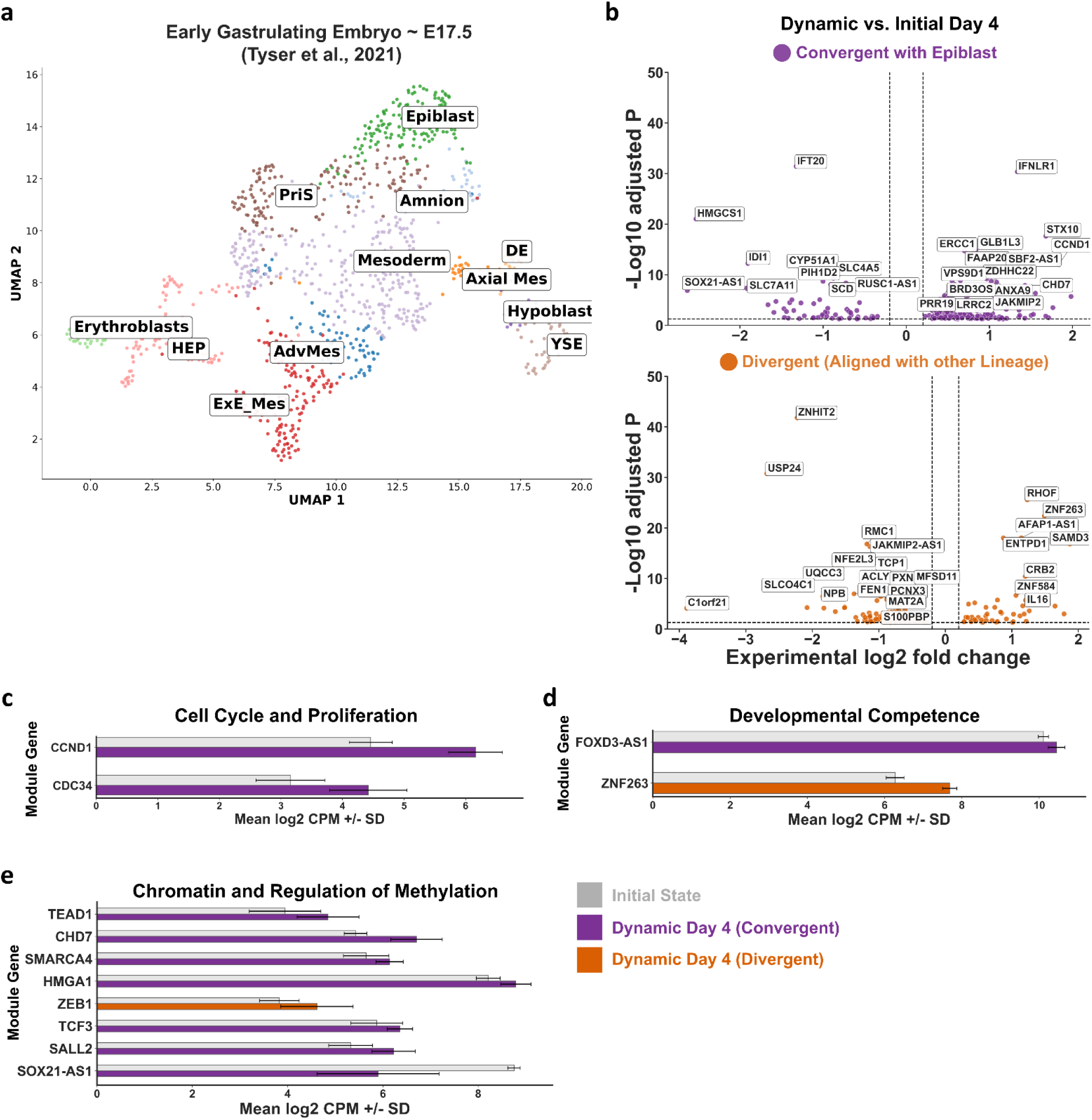
(a) UMAP projection of ∼E17.5 early gastrulating embryonic cell types (Tyser et al., 2021). Exe_Mes: extraembryonic mesoderm; AdvMes: advanced mesoderm; YSE: yolk salk endoderm; Axial Mes: axial mesoderm; DE: definitive endoderm; HEP: hemato-endothelial progenitors; PriS: primitive streak. (b) Volcano plots visualizing the number of dynamic vs. initial differentially expressed genes that converge on epiblast or diverge towards downstream lineages. Convergence within gene modules of interest in dynamic vs. initial are plotted for (c) cell cycle and proliferation, (d) developmental competence, and (e) chromatin and regulation of methylation genes.

Genes significantly altered in both an experimental contrast and an Epiblast-versus-lineage contrast were classified according to directional alignment. Genes changing in the same direction as the corresponding epiblast-versus-lineage effect were designated epiblast-aligned, or convergent. Genes changing in the opposing direction were designated lineage-aligned, or divergent.

Transition from static aggregates to dynamic suspension produced a broad transcriptional intersection with the developmental reference. The dynamic response included both Epiblast-aligned and lineage-aligned genes across the evaluated embryonic and extraembryonic contrasts, with an overall predominance of Epiblast-aligned relationships (Supplementary Figure 4a). This pattern indicates that adaptation to dynamic culture heterogeneously organizes genes implicated in both epiblast and other-lineage identities, independent of core pluripotency readouts.

Differential analysis separated a subset of overall differential dynamic response into epiblast-aligned and lineage-aligned components (Figure 5b). Epiblast-aligned genes included regulators associated with lipid and sterol metabolism, cell-cycle control, chromatin regulation, genome maintenance, and developmental competence. The convergent component included genes such as HMGCS1, IDI1, SOX21-AS1, ERCC1, BRD4, CHD7, and CCND1. In contrast, the lineage-aligned component included genes associated with transcriptional regulation, RNA processing, cellular architecture, and developmental identity, including ZNHIT2, ZNF263, RHOF, NFE2L3, and PXN. Cell-cycle and proliferation genes included epiblast-aligned increases in CCND1 and CDC34 (Figure 5c). Developmental competence was similarly divided, with FOXD3-AS1 showing Epiblast alignment and ZNF263 showing alignment with non-pluripotent lineages (Figure 5d). Chromatin and transcriptional regulators, including CHD7, SMARCA4, HMGA1, TCF3, SALL2, and SOX21-AS1, were predominantly epiblast-aligned, whereas ZEB1 showed non-pluripotent lineage alignment (Figure 5e).

Increasing agitation from 120 to 150 RPM produced only three genes implicated in differential identity across early gastrulating embryo. Jointly differential genes were detected across several epiblast-versus-lineage contrasts, but the number of overlapping genes was limited to only 3 genes (Supplementary Figure 4h). These three genes captured the entire agitation-associated intersection: KCNK12, LARP7, and MMP24OS (Supplementary Figure 4i). LARP7 and MMP24OS showed epiblast-alignment, whereas KCNK12 showed a lineage-alignment. These genes implicate membrane excitability, RNA-associated regulation, and transcription linked to the MMP24 locus as distinct components of the agitation response. However, the small number of jointly differential genes does not support a coherent pathway- or module-level interpretation.

## Discussion

This study demonstrates that hiPSC expansion conditions can preserve core pluripotency-associated features while substantially reorganizing metabolic, transcriptional, and proteomic state within a single passage. Culture configuration altered translation-associated programs, sterol metabolism, mitochondrial organization, mechanotransduction, and proteins surrounding the pluripotency network despite the stability of OCT4, SOX2, and NANOG [40, 41]. Differences in cotranslational targeting, elongation, and translation-associated programs are also consequential because translational control may contribute to stem-cell maintenance and cell-fate reorganization [42]. These findings indicate that conventional marker retention may not establish equivalence of developmental competence among expanded populations.

Differences in DPPA4, DNMT3B, and FGF2 are relevant even without corresponding loss of OCT4, SOX2, or NANOG. DPPA2 and DPPA4 have been implicated in maintaining bivalent developmental promoters, protecting developmental loci from DNA methylation, and preserving competence for subsequent differentiation [43]. DPPA4 can also be dispensable for maintenance of embryonic stem-cell identity while remaining important for later embryonic development, illustrating why conventional pluripotency-marker retention and developmental competence should not be treated as equivalent [44]. The present data do not demonstrate altered chromatin accessibility or DNA methylation directly. Rather, the observed differences identify culture-dependent variation in molecular regulators capable of influencing how an expanded population subsequently organizes in response to differentiation signals.

Transition into dynamic suspension produced the broadest molecular response. Dynamic culture was associated with mitochondrial reorganization, altered carbon and lipid allocation, changes in proteostasis and RNA regulation, and remodeling of adhesion, cytoskeletal, cell-cycle, chromatin-associated, and developmental programs. Adhesion, cytoskeletal tension, matrix interaction, and externally applied forces can influence intracellular signaling, while metabolic pathways can affect transcription and the epigenetic landscape through metabolite availability, mitochondrial signaling, and direct co-factor engagement [45, 46]. The burst of differential metabolic and transcriptional activity observed by day 2 under normoxia relative to hypoxia is suggestive of a metabolic state remodeling to support adaptation to the environment; the lack of differential transcriptional activity at day 4 coupled with remodeling of the metabolic and energetic adaptation transcriptome under dynamic conditions generally could suggest competing compensatory and adaptive programs that are difficult to decouple in single-passage observations.

The reciprocal differences in tryptophan and kynurenine provide an additional metabolic dimension to the response. Tryptophan metabolism is implicated in human pluripotent stem-cell proliferation and maintenance, while kynurenine-pathway activity differs among pluripotent states and connects tryptophan utilization with redox regulation, bioactive signaling metabolites, and de novo NAD synthesis [47]. Oxygen-dependent redistribution of these metabolites may reflect altered allocation of tryptophan between proliferation-associated metabolism and downstream kynurenine-pathway activity. This finding does not establish increased pathway flux, AhR activation, or assess downstream lineage, but it identifies plausible activity through which oxygen availability might alter metabolic and regulatory state despite preservation of core pluripotency markers and normalization of carbon bioenergetic throughput.

The dynamic environment establishes a heterogeneous adaptive state in which selected metabolic, structural, and regulatory programs align either with both epiblast and downstream developmental lineages. This interpretation is consistent with evidence that mechanical and metabolic cues act together to regulate pluripotent-cell state. Transcriptionally, ZNF263 is notable because ZNF263 has been shown to initiate early differentiation-gene expression, dampen core pluripotency circuitry, and facilitate pluripotency dissolution and primary germ-layer commitment, making its alignment a plausible indicator of increased lineage priming [32]. NFE2L3 adds a transcriptional stress-adaptation component, as NFE2L3 has been linked to differentiation, cellular stress responses, and inflammatory regulation [48]. Dynamic culture establishes a heterogeneous, developmentally poised state in which epiblast-associated proliferation and regulatory competence coexist with selective activation of transcriptional, adhesive, cytoskeletal, and RNA-regulatory machinery associated with lineage responsiveness. Further, while the predominance of epiblast-aligned genes supports retention of a primed pluripotent identity, the simultaneous presence of lineage-aligned gene expression indicates that conventional pluripotency-marker stability does not establish molecular equivalence with nature.

### Limitations

There are a number of limitations to this work. The most interesting limitation is the lack of epigenomics data in this work. A major unresolved question is whether longitudinal cell state compensation to perturbation leaves lasting effects on 3-dimensional chromatin architecture or accessibility of genes. Secondly, the breadth and longitudinal scope of this work resulted in a large number of samples across the set; however statistically future work would benefit from n > 5, 7, or even 10 biological replicates, particularly given the variance that is observed across biomanufacturing batches in maintaining and propagating hiPSCs, even under well-defined protocols. It is also pertinent to scale the number of cell lines utilized. The effect of reprogramming strategy on these nuances is entirely unknown, and no definitive argument can be made as to the generalizability of our current findings. Importantly, this work focused on single-passage dynamics. Additional studies are needed to assess the impact of phenotypic reorganization across the continuum of expansion and derivation of down-stream derivatives.

## Materials and Methods

### hiPSC Culture Protocols

All procedures were approved by the Conjoint Health Research Ethics Board (CHREB) at the University of Calgary under ethics protocol #REB14-1914.

### Proteomics cultures

Experiments were carried out using the hiPSC line TC1133 (Lonza, USA). For the seed train, cells were thawed and expanded through two passages on Matrigel-coated T-75 flasks, seeded at 15,000 cells/cm² for 72 h followed by 5,000 cells/cm² for 88 h. Cultures were maintained in mTeSR1 medium at 37°C and 5% CO₂, with daily 100% medium exchanges at 0.2 mL/cm². Reagents were supplemented with 10 µM Y-27632 during thawing and passaging. Passaging was performed using Accutase and mTeSR1, both supplemented with Y-27632. Following seed train expansion, cells were inoculated at 20,000 cells/mL into T-25 flasks (static adherent), 6-well suspension plates (suspended aggregates), or PBS-0.1 Mini Vertical-Wheel bioreactors (agitated aggregates). Static and suspension cultures were maintained at 0.2 mL/cm², while bioreactors were operated at a 100 mL working volume. To assess hydrodynamic effects, bioreactors were agitated at 20 RPM (below range), 60 RPM (within range), or 100 RPM (above range). All experimental cultures were maintained in mTeSR1 medium without Y-27632. T-25 cultures received daily 100% medium exchanges. Suspension plates and bioreactors received a single 50% exchange on Day 3. Four biological replicates per condition were harvested on Day 4 for processing and proteomic analysis.

### Metabolomics and transcriptomics cultures

Cells from the hiPSC line PGPC14 (Ellis Lab, University of Toronto) were cultured in 18.5 mL parallel laboratory-scale bioreactors and characterized as previously reported [23]. An inoculation density of 1×10^4^ cells/cm^2^ was used for aggregate preformation. All cell populations were cultured in mTeSR1, with 10 µM Y-27632 supplemented for aggregate preformation and dynamic culture. All dynamic cultures were inoculated at 3.6×10^4^ cells/mL. Cultures were maintained at 37 °C, greater than 88% relative humidity, and 5% CO_2_. Atmospheric oxygen was maintained at approximately 19.7% O_2_ for normoxic cultures or 5% O_2_ for hypoxic cultures. Chamber oxygen was monitored and controlled using a ProOx P110 compact O_2_ controller with an external probe, while CO_2_ was regulated using a ProCO_2_ Model 120 controller and external probe (BioSpherix, New York, USA). An external 1 liter water reservoir was maintained within the incubator to limit evaporation. Control samples were taken following aggregate preformation, denoted as initial state. Dynamic samples were taken at days 2 and 4. Two technical replicates were obtained from each bioreactor and pooled, from three separate biological replicates for all conditions at all timepoints. All sample volumes were 1 mL with replacement.

## Omics

### Proteomics Sample Preparation

Cell samples were lysed in 500 µL buffer (2% SDS, 200 mM ammonium bicarbonate, protease inhibitors) with 4–5 stainless-steel beads, homogenized at 30 Hz for 20 min and sonicated (Fisherbrand™ Model 120). Lysates were centrifuged at 14,000 x g at 4 °C, and supernatants collected in Protein LoBind tubes. Protein concentrations were measured using ThermoFisher BCA assay. For each sample, 100 µg protein was TCA-precipitated, incubated on ice for 30 min, centrifuged at 14,000 x g for 15 min at 4 °C, washed three times with ice-cold acetone, and stored at −20 °C overnight. Pellets were resuspended in 8 M urea/Tris, reduced with 10 mM DTT at 37 °C for 30 min, and alkylated with 50 mM iodoacetamide in the dark for 25 min. Samples were loaded onto 30-kDa filters, centrifuged, and washed three times with 8 M urea/Tris and three times with 50 mM ammonium bicarbonate. Proteins were digested overnight at 37 °C with trypsin (1:10), and peptides were eluted with 50 mM ammonium bicarbonate. Peptides were cleaned using Waters C18 SPE columns following the manufacturer instructions.

Tryptic peptides were analyzed using an Orbitrap Fusion Lumos Tribrid mass spectrometer (Thermo Scientific) with Xcalibur v4.4.16.14, coupled to an Easy-nLC 1200. A total of 2 μg peptide was loaded onto a C18 trap (75 μm x 2 cm, PepMap 100; P/N 164946) at 2 μL/min in solvent A (0.1% formic acid in water), then separated on a C18 analytical column (75 μm x 50 cm, PepMap RSLC; P/N ES803) at 0.3 μL/min. Peptides were eluted using a 120-min gradient from 5 - 40% solvent B (0.1% formic acid in 80% acetonitrile). Electrospray ionization was performed at 2.1 kV into a 300°C ion transfer tube in positive mode. Full MS scans were acquired in the Orbitrap at 120,000 FWHM over m/z 375–1575 (charges +2 to +7) with 4 ×10^5^ AGC and 50 ms injection time. The instrument operated in top-speed mode with a 3 s cycle. Precursors ≥ 5000 intensity with peptide-like isotopic profiles were quadrupole-isolated and fragmented by HCD (30% CE). MS2 spectra were collected in the ion trap at a rapid scan rate (AGC 1×10^4^; 35 ms injection time). Dynamic exclusion was set to 45 s with a ± 10 ppm tolerance.

### Metabolomics Sample Preparation

Protocols were carried out as previously described [49, 50]. Solutions of 90% and 50% HPLC-grade methanol in culture-grade water were prepared and stored at - 20°C prior to immediate use. Samples taken from culture were immediately centrifuged at 500xg for 5 minutes. Following centrifugation, the supernatant was aspirated, and cells were resuspended in 2 mL 90% methanol. Samples were refrigerated for 10 minutes on ice. The solution was centrifuged at 4000xg for 15 minutes at 4°C. Supernatant was isolated and centrifuged at 4000xg under vacuum for 24h at 4°C. Samples were resuspended in 100 µL of 50% methanol and stored at -80°C prior to analysis at the Calgary Metabolomics Research Facility (CMRF). Semi-targeted metabolic analysis was performed on a Q Exactive Hybrid Quadrupole-Orbitrap Mass Spectrometer coupled to a Vanquish UHPLC System (Thermo-Fisher, USA). Chromatographic separation of metabolites was performed on a Zorbax SB-C18 UHPLC column at a flow rate of 600 µL/min (Agilent, Cat# 822700-902). A binary solvent system was used: Solvent A, 10 mM tributylamine, 10 mM acetate pH 7.5 in 97/3% (v/v) mass spectrometry grade water/methanol, and solvent B, MS grade acetonitrile. A sample injection volume of 2 µL was used. The spectrometer was operated in negative full scan mode with a resolution of 140,000 scanning from 50-750 m/z. Spectroscopy data was processed in EI-MAVEN. Metabolites were identified by matching m/z signals (±10 ppm) and chromatographic retention times relative to LMSLS commercial standards (Sigma-Aldrich).

### Transcriptomics Sample Preparation

Samples taken from culture were immediately centrifuged at 500xg for 5 minutes. Following centrifugation, supernatant was aspirated and cells were resuspended in 2.5 mL RNAlater stabilization solution (Thermo-Fisher, Cat# AM7023). These samples were stored at 4°C for 24 hours, then transferred to -80°C storage until ready for RNA processing and sequencing. Samples were thawed at ambient temperature and purified using a QIAGEN RNeasy Mini Kit (Cat# 74104) following the manufacturers protocols. The final samples were adjusted for target concentrations of 50 ng/µL in 15 µL of RNAse free water (Thermo-Fisher, Cat# 10977015) and submitted to the Centre for Health Genomics and Informatics (CHGI) at the University of Calgary for analysis. RNA library preparation was carried out with a NEBNext Ultra II Directional RNA Library Prep Kit using the manufacturer’s protocols (Illumina, Cat# E7760L). Total RNA sequencing with ribosomal depletion and preparation of fastq files was carried out using a NovaSeq 6000 and DRAGEN Bio-IT platform (Illumina, USA).

## Data Processing

### Proteomics

Mass spectrometry spectra were searched against the *Homo sapiens* UniProt reference proteome FASTA, downloaded February 24, 2022, using MaxQuant v1.6.0.1. Default MaxQuant settings were used except that oxidation of methionine, protein N-terminal acetylation, and deamidation of asparagine or glutamine were included as variable modifications. Label-free quantification was enabled for the agitation experiment. The first-search peptide tolerance was 10 ppm, the FASTA identifier rule was set to UniProt, the maximum peptide mass was 6,600 Da, the minimum peptide length was five amino acids, and match between runs was enabled. Peptide-spectrum matches and protein identifications were controlled at a false-discovery rate below 1%.

Reverse-sequence identifications, potential contaminants, and proteins identified only by modification site were excluded before statistical analysis. Gene symbols were assigned from the MaxQuant protein-group annotations. The culture-platform experiment consisted of three pairwise SILAC comparisons: static aggregate versus dynamic aggregate, static monolayer versus dynamic aggregate, and static aggregate versus static monolayer. Replicate-level normalized heavy-to-light ratios were log2 transformed and analyzed jointly using a connected linear model. The model estimated centered relative protein abundance for static monolayer, static aggregate, and dynamic aggregate culture while preserving the orientation of each SILAC comparison. Platform-specific effects were calculated by comparing each platform with the mean of the other two platforms. Pairwise platform contrasts were also calculated directly from the connected model.

For the agitation experiment, MaxQuant LFQ intensities were log2 transformed and analyzed using a three-group linear model comparing cultures maintained at 20, 60, or 100 RPM. Pairwise contrasts were calculated for 60 versus 20 RPM, 100 versus 60 RPM, and 100 versus 20 RPM. Agitation-specific effects were calculated by comparing each RPM condition with the mean of the other two conditions.

For both experiments, residual protein-level variances were stabilized using empirical-Bayes moderation. Moderated t-statistics and two-sided p-values were calculated for each contrast, followed by Benjamini-Hochberg correction within each contrast. Proteins were classified as differentially abundant using an adjusted p-value below 0.05 and an absolute log2 fold change of at least log2(1.5).

Pathway analysis was performed by pre-ranked gene-set enrichment analysis using moderated t-statistics as the ranking metric. Gene sets were obtained from MSigDB Hallmark 2020, Reactome 2022, and Gene Ontology Biological Process 2023 collections. Enrichment was calculated using 1,000 permutations, with gene-set sizes restricted to 10 – 500 genes. Normalized enrichment scores and false-discovery rates were calculated separately for each contrast. Pathways were selected using their enrichment magnitude and false-discovery rate and hierarchically clustered by normalized enrichment-score profiles using Euclidean distance and average linkage. All statistical analyses and visualizations were generated using versioned Python workflows, with source tables exported for each protein-level contrast, pathway analysis, and figure panel.

### Metabolomics

Intracellular metabolite peak intensities were corrected by subtracting the median intensity measured across blank injections. Negative values following blank subtraction were set to zero. Metabolites were retained when detected above the corrected baseline in at least 70% of biological samples and when the median biological-sample intensity was at least threefold greater than the median blank intensity. Retained metabolite abundances were normalized using probabilistic quotient normalization and log2 transformed before statistical analysis. Metabolite names were standardized across the dataset, and blank, calibration-standard, and control injections were excluded from biological comparisons.

Differential metabolite abundance was assessed using two-group linear models with empirical-Bayes moderation of metabolite-level residual variances. Comparisons included normoxia versus hypoxia, pairwise agitation contrasts among 100, 120, and 150 RPM, and oxygen- or agitation-specific contrasts within the relevant experimental strata. For analyses pooled across days 2 and 4, repeated measurements from the same independently operated bioreactor were averaged to generate one subject-level value, preventing repeated timepoints from being treated as independent biological replicates. Day-specific comparisons used the biological replicates collected at the corresponding timepoint.

Moderated t-statistics and two-sided p-values were calculated for each metabolite. P-values were adjusted separately within each contrast using the Benjamini-Hochberg procedure. Differentially abundant metabolites were defined by an adjusted p-value below 0.05 and an absolute log2 fold change of at least log2(1.5). Complete results, including effect sizes, moderated standard errors, raw p-values, adjusted p-values, and group sample sizes, were retained for all tested metabolites.

Condition-level heatmaps were generated from mean log2-normalized metabolite abundance and displayed as metabolite-wise Z scores. Metabolites were ordered by average-linkage hierarchical clustering using Euclidean distance. For comparison-specific heatmaps, all metabolites meeting the false-discovery threshold were retained, followed by the highest-ranking remaining metabolites according to the absolute moderated t-statistic. Volcano plots display log2 fold changes against negative log10-adjusted p-values. All preprocessing, statistical analyses, and visualizations were performed using versioned Python workflows, with normalized abundance matrices, sample metadata, quality-control summaries, statistical results, and figure-source tables exported for reproducibility.

### Transcriptomics

Adapter-trimmed FASTQ files were processed using an offline RNA-sequencing workflow. Sequence quality was assessed using FastQC v0.12.1, and reads were aligned to the human GRCh38 reference genome using HISAT2 v2.2.1. Alignment and sequencing quality metrics were consolidated using MultiQC v1.11. Gene-level count files were generated from the aligned reads and imported into a versioned Python analysis workflow. Technical libraries and sequencing lanes corresponding to the same biological sample were summed before statistical analysis. Genes were retained when they had at least 10 counts in a minimum of three biological samples.

Differential expression was assessed from raw integer counts using PyDESeq2, with Benjamini-Hochberg correction applied independently within each contrast. Genes were classified as moderately differentially expressed when the adjusted p-value was below 0.05 and the absolute log2 fold change was at least 0.2, and as large-effect differentially expressed when the adjusted p-value was below 0.05 and the absolute log2 fold change was at least log2(1.5). For comparisons pooled across days 2 and 4, repeated measurements from each independently operated bioreactor were summed to create one subject-level count profile, preventing longitudinal samples from being treated as independent biological replicates. Day-specific comparisons retained the biological replicates collected at corresponding timepoints. Complete differential-expression results, including mean abundance, log2 fold change, standard error, Wald statistic, raw p-value, and adjusted p-value were retained for every tested gene.

For expression visualization, raw counts were converted to counts per million and log2 transformed after addition of a pseudocount. Condition-level heatmaps were generated from biological-sample means and displayed as gene-wise Z scores. Gene-level bar plots show mean log2 counts per million and standard deviation across biological samples. Over-representation analysis was performed using differentially expressed genes against the set of genes tested in the corresponding contrast as the statistical background. Gene Ontology Biological Process terms were tested separately using coupled, upregulated, and downregulated DEG sets, with Benjamini-Hochberg correction applied to enrichment p-values. Volcano plots display log2 fold changes against negative log10-adjusted p-values. All statistical analyses and visualizations were performed using versioned Python workflows, with count matrices, sample metadata, quality-control summaries, complete contrast tables, pathway results, and figure-source data exported for reproducibility and stored under the GitHub repository for this work.

### Developmental Reference and Cross-dataset Analysis

Publicly available single-cell RNA-sequencing counts and metadata were obtained from the Tyser et al. early human gastrulation dataset E-MTAB-9388 [41]. The reference dataset was processed independently from the experimental transcriptomic data and was not integrated with the experimental count matrix. Cells with ambiguous annotations or prediction confidence below 0.50 were excluded from developmental-state comparisons. The predicted annotation was used as the primary cell-state label, while the original and fine-grained annotations were retained for provenance. Analyses focused on Epiblast, primitive streak, mesoderm, and definitive endoderm.

Raw reference counts were normalized to counts per million and transformed as log1p(CPM). Differential expression was calculated between Epiblast and each alternative developmental state using two-sided Mann-Whitney U tests on log1p(CPM) values. Benjamini-Hochberg correction was applied independently within each pairwise comparison. Experimental and reference results were matched by standardized gene symbol. Cross-dataset analyses were restricted to genes meeting the moderate differential-expression threshold in both the experimental contrast and the corresponding developmental reference contrast (Padj < 0.05, Log2FC > 0.2). Significant genes with experimental and reference log2 fold changes of the sign were classified as Epiblast-aligned or convergent. Those with opposing signs were classified as lineage-aligned or divergent. Because each reference contrast was oriented as Epiblast versus developmental state, convergence indicated alignment with the Epiblast side of the contrast, whereas divergence indicated alignment with primitive-streak, mesodermal, or definitive-endoderm (Eq. 1 – 3).

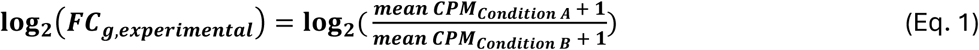

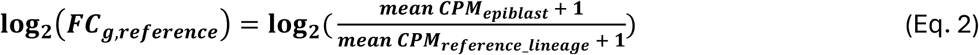

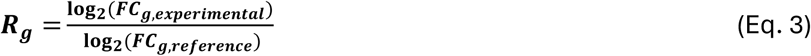

For pooled developmental analyses, genes were classified as epiblast-aligned when they were convergent in at least one developmental comparison and never divergent, or as lineage-aligned when they were divergent in at least one comparison and never convergent. Genes showing different directional classifications across primitive streak, mesoderm, and definitive endoderm were designated as mixed and were not included in pooled convergent or divergent enrichment analyses.

Over-representation analysis was performed separately for epiblast-aligned and lineage-aligned gene sets using the genes tested in the corresponding experimental contrast as the statistical background. Gene Ontology Biological Process terms were evaluated using a hypergeometric test, followed by Benjamini-Hochberg correction. Cross-dataset analyses were performed for dynamic versus initial culture, 150 versus 100 RPM, and normoxia versus hypoxia. Results were summarized using directional gene counts, effect-size heatmaps, experimental volcano plots, normalized expression bar plots, and alignment-specific pathway enrichment.

## Author Contributions

J.C. conceived the project, designed and carried out all metabolomics and transcriptomics experiments, wrote software, analyzed the multi omics data, wrote the manuscript, and prepared all Figures. T.D. designed and carried out proteomics experiments, analyzed cell expansion data, supported figure preparation, and edited the manuscript. E.G. contributed to proteomics experimental design, sampling, and manuscript edits. D.Y. carried out proteomics protocols. A.D. edited the manuscript. I.L., M.K. and K.M. acquired funding for this work, reviewed data, challenged findings, provided mentorship, contributed intellectually, and edited the manuscript.

## Data Availability

Proteomics data from this work can be accessed from PRIDE (PXD078823). Metabolomics data from this work can be accessed from Metabolomics Workbench (ST004863). Bulk RNASeq data from this work can be accessed from GEO (GSE331399). Reference data for early gastrulating embryo (E-MTAB-9388) was obtained from GEO [46]. All extended data, including reference counts, tables, and scripts for all analytical work is hosted on GitHub (https://github.com/JxColter/ColterJ_etal_Omics). Any additional material is available upon request from the corresponding author.

## Acknowledgements

The authors would like to acknowledge Dr. Marija Drikic and Dr. Annegret Ulke-Lemee at the Calgary Metabolomics Research Facility (CMRF). The authors further acknowledge that acquisition of metabolomics data at the CMRF was supported by the International Microbiome Centre and the Canada Foundation for Innovation (CFI-JELF 34986). The authors would also like to acknowledge Shelly Wegener, Dr. Paul Gordon, Jené Weatherhead, Juli Kriel, and Caroline Quesnel at the Calgary Centre for Health Genomics and Informatics (CHGI) for their contributions to acquisition and preprocessing of transcriptomics data. Lastly, the authors gratefully acknowledge Dr. James Ellis’ lab for providing one of the cell lines utilized in the second study.

