## Supplementary Figures for "Multi-omics analyses reveal environmentally adaptive cell-state remodeling during human iPSC biomanufacturing"

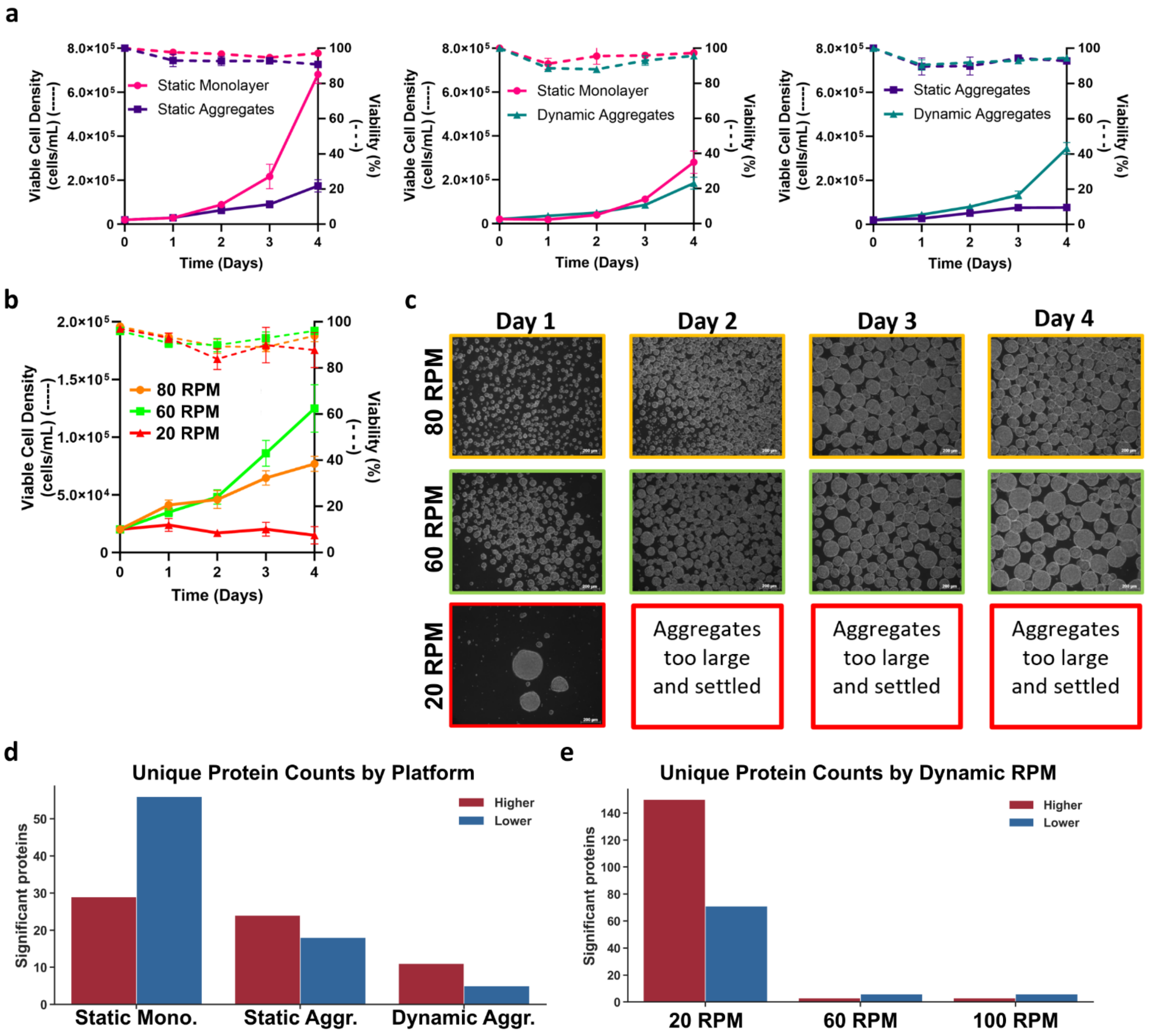


**Supplementary Figure 1:** (a) Growth kinetics for comparisons of static monolayer, static aggregates, and dynamic aggregates. (b) Growth kinetics under dynamic conditions. (c) Morphological characterization under dynamic culture from days 1 to 4. Counts for differentially significant protein abundances are shown for (d) platforms and (e) dynamic cultures by RPM.

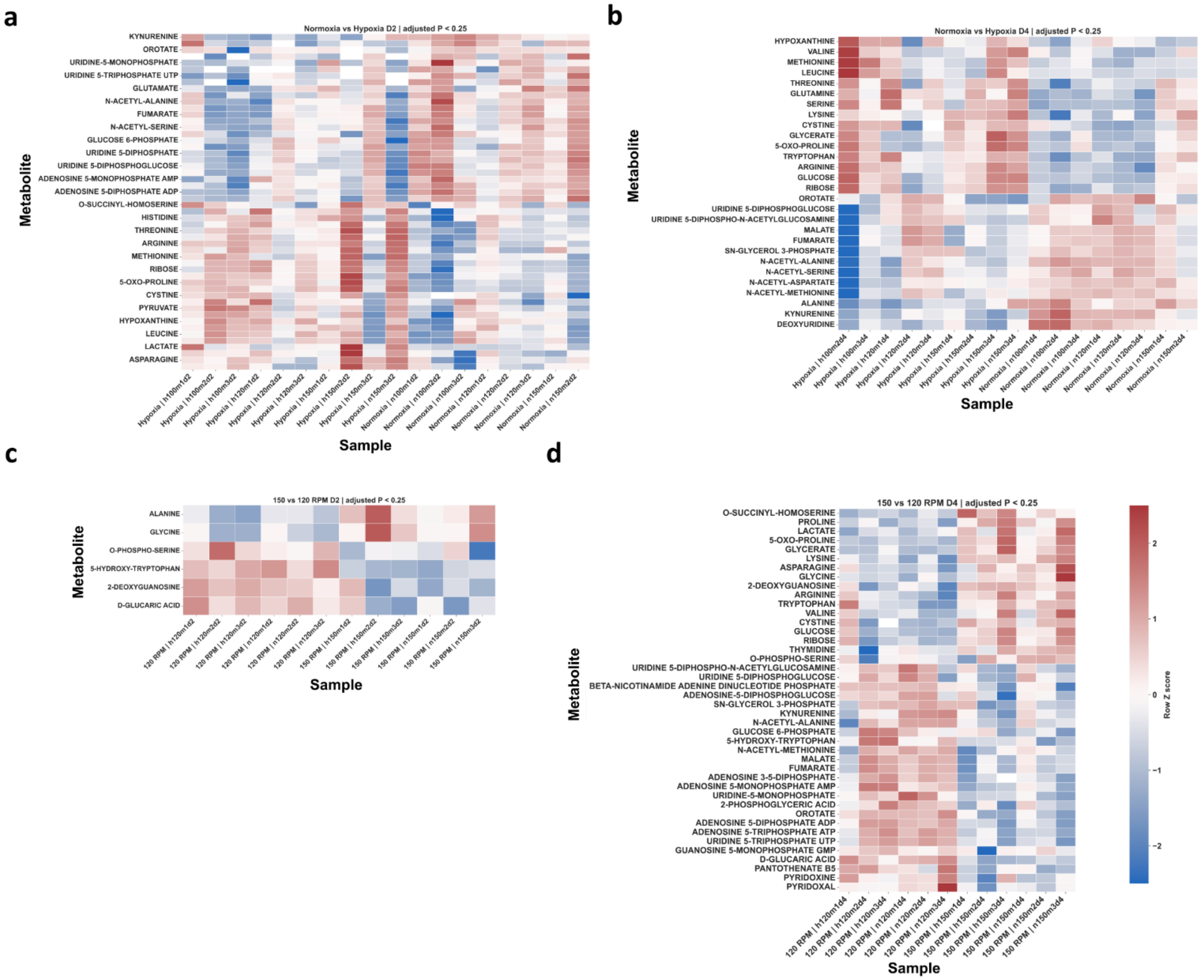


**Supplementary Figure 2:** Normalized z scores of metabolic abundances, organized by sample for normoxia and hypoxia at (a) day 2 and (b) day 4, and for 120 RPM and 150 RPM at (c) day 2 and (d) day 4.

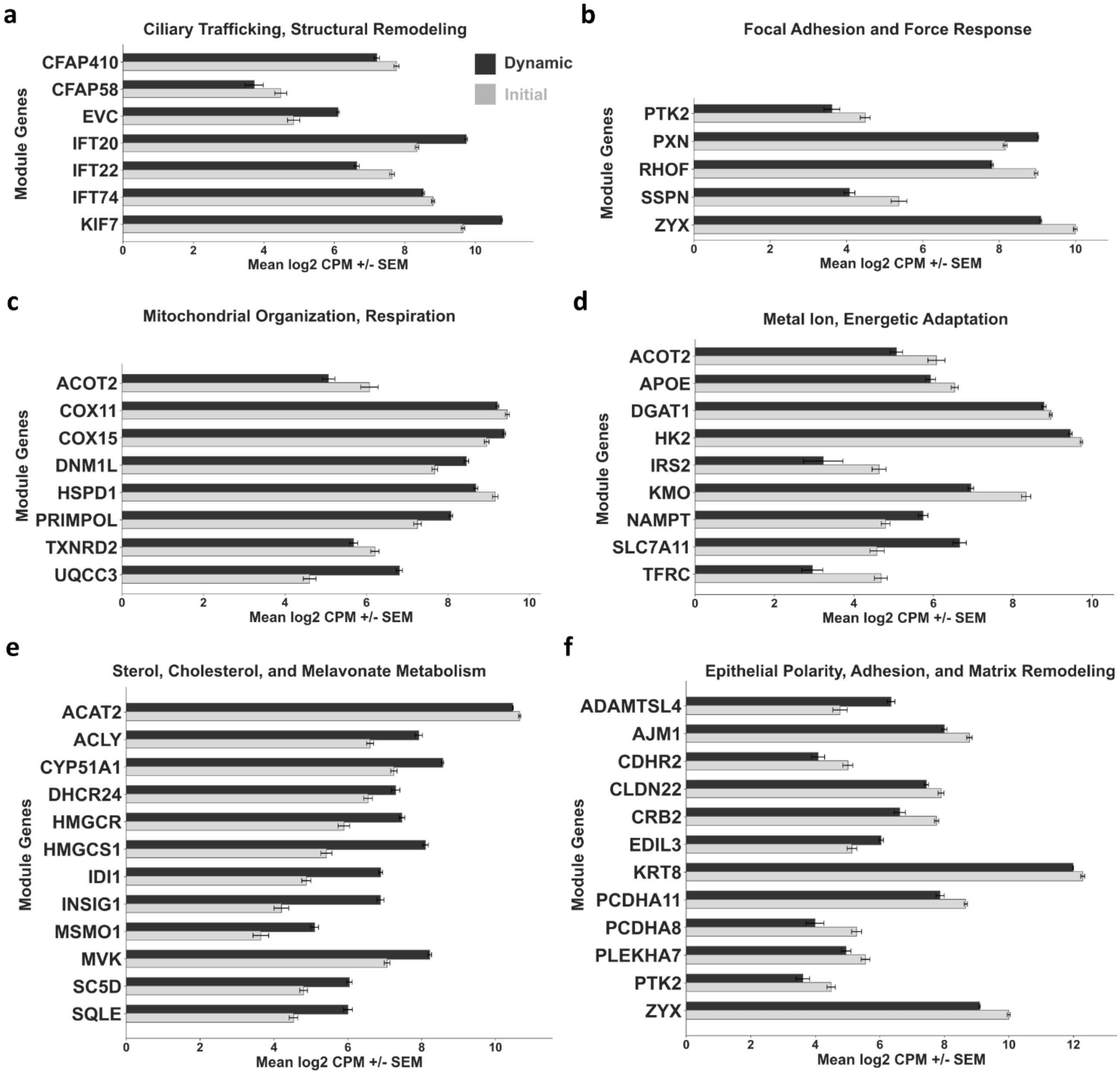


**Supplementary Figure 3:** Significant DEGs for curated gene modules for (a) ciliary trafficking, structural remodeling, (b) focal adhesion and force response, (c) mitochondrial organization, respiration, (d) metal ion, energetic adaptation, (e) sterol, cholesterol, and mevalonate metabolism, and (f) epithelial polarity, adhesion, and matrix remodeling.

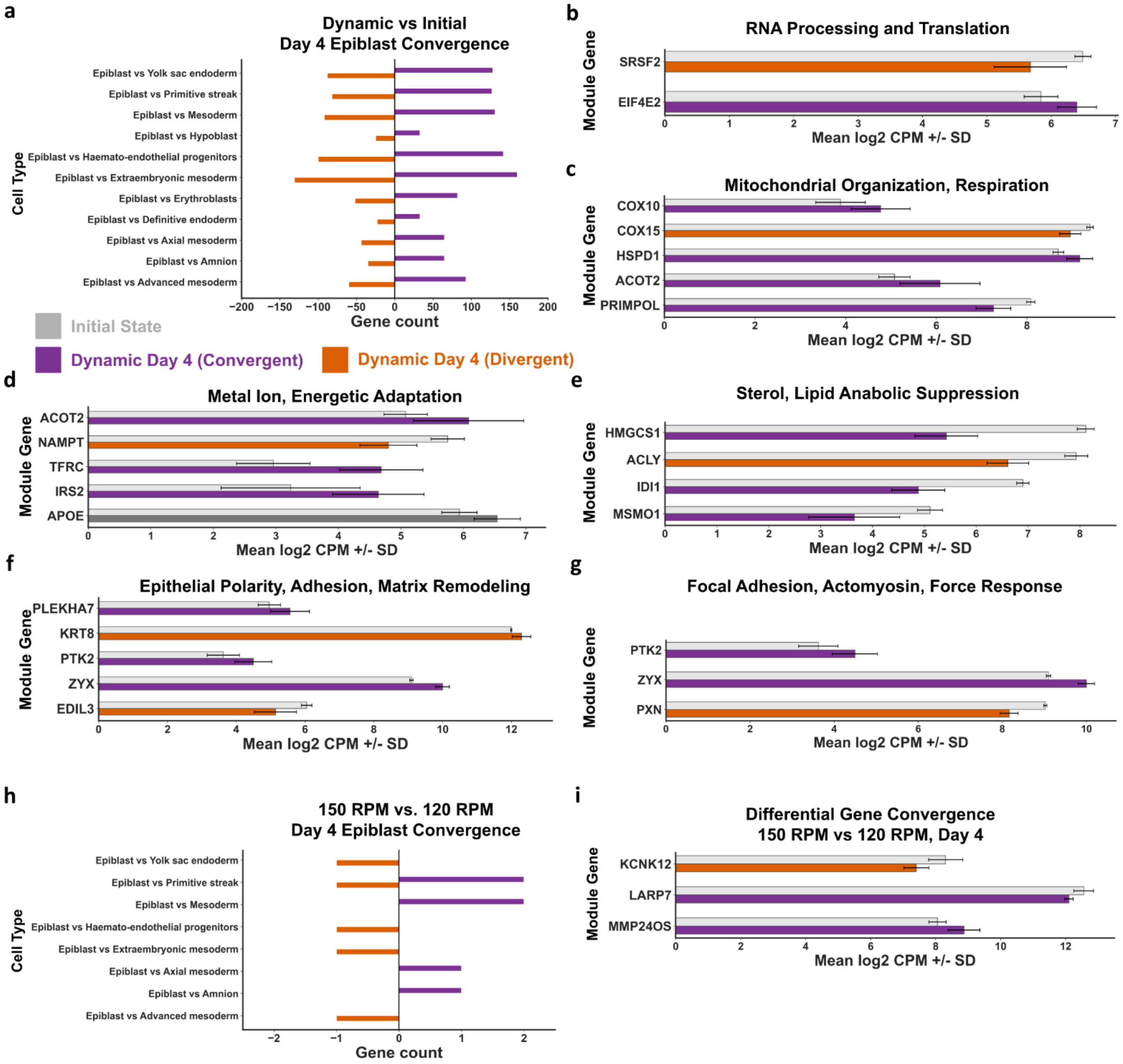


**Supplementary Figure 4:** (a) Convergence with epiblast for dynamic vs. initial state against non-pluripotent lineages. Significant convergent and divergent DEGs under dynamic vs. initial state for curated gene modules for (b) RNA processing and translation, (c) mitochondrial organization, respiration, (d) metal ion, energetic adaptation, (e) sterol, lipid anabolic suppression, (f) epithelial polarity, adhesion, and matrix remodeling, and (g) focal adhesion and force response. (h) Convergence with epiblast for 150 RPM vs. 120 RPM against non-pluripotent lineages. (i) DEGs identified to fall within those differentially expressed in early gastrulation.
